# Modeling and predicting single-cell multi-gene perturbation responses with scLAMBDA

**DOI:** 10.1101/2024.12.04.626878

**Authors:** Gefei Wang, Tianyu Liu, Jia Zhao, Youshu Cheng, Hongyu Zhao

## Abstract

Understanding cellular responses to genetic perturbations is essential for understanding gene regulation and phenotype formation. While high-throughput single-cell RNA-sequencing has facilitated detailed profiling of heterogeneous transcriptional responses to perturbations at the single-cell level, there remains a pressing need for computational models that can decode the mechanisms driving these responses and accurately predict outcomes to prioritize target genes for experimental design. Here, we present scLAMBDA, a deep generative learning framework designed to model and predict single-cell transcriptional responses to genetic perturbations, including single-gene and combinatorial multi-gene perturbations. By leveraging gene embeddings derived from large language models, scLAMBDA effectively integrates prior biological knowledge and disentangles basal cell states from perturbation-specific salient representations. Through comprehensive evaluations on multiple single-cell CRISPR Perturb-seq datasets, scLAMBDA consistently outperformed state-of-the-art methods in predicting perturbation outcomes, achieving higher prediction accuracy. Notably, scLAMBDA demonstrated robust generalization to unseen target genes and perturbations, and its predictions captured both average expression changes and the heterogeneity of single-cell responses. Furthermore, its predictions enable diverse downstream analyses, including the identification of differentially expressed genes and the exploration of genetic interactions, demonstrating its utility and versatility.

## Introduction

Understanding how cells respond to specific perturbations is a fundamental question in biology, as it provides critical insights into the mechanisms that govern cellular behavior [1]. Recent advances in integrating clustered regularly interspaced short palindromic repeats (CRISPR) gene-editing technology with high-throughput single-cell RNA-sequencing (scRNA-seq) approaches, such as Perturb-seq [2], enable detailed profiling of transcriptional responses in individual cells subjected to genetic perturbations. These tools provide critical insights into complex gene regulatory networks and the role of gene expression in shaping phenotypes [3], with promising applications in clinical trials and targeted therapies [4, 5, 6].

Despite these advancements, conducting large-scale genetic perturbation screens for numerous target genes and their combinations remains a labor-intensive and resource-demanding endeavor [7]. Furthermore, single-cell molecular profiling typically destroys cells, complicating the analysis of perturbation mechanisms with potential cell heterogeneity. There is a great need for computational approaches that can elucidate the mechanisms underlying perturbations, predict outcomes, and guide experimental designs efficiently [8].

Many computational tools have been developed to understand the causality and heterogeneity behind single-cell perturbation effects using factor analysis or deep generative modeling approaches, such as GSFA [9], contrastiveVI [10], sparse VAE [11] and SAMS-VAE [12]. However, these models are not specifically designed to predict transcriptional responses to perturbations. In contrast, several methods have been developed to predict perturbation effects. For instance, scGen [13] and CellOT [14] facilitate *in silico* prediction of cellular responses to previously observed types of perturbations, while CPA [15] extends this to combinations of observed perturbations. Despite their strengths, these approaches are constrained by their reliance on training data that include either single-gene perturbations or all target genes involved in multi-gene perturbations, which limits their applicability for predicting outcomes for novel target genes. Recently, several methods including GEARS [16], GenePert [17], and single-cell foundation models such as scGPT [18] have been developed for predicting the perturbation effects of novel unseen genes. However, GEARS and GenePert focus primarily on average perturbation effects, not able to capture single-cell level heterogeneity. Additionally, foundation models for perturbation prediction have yet to demonstrate consistently better performance than simple baselines [19, 20].

To address the limitations of existing computational methods, we present scLAMBDA, a deep generative model for predicting single-cell genetic perturbation responses with LAnguage Model-assisted emBedding-Disentangled Autoencoders. By leveraging the embedding capabilities of large language models [21, 22, 23], scLAMBDA effectively predicts genetic perturbation outcomes for unobserved target genes or gene combinations. Its disentangled representation learning framework enables the modeling of single-cell level perturbations by separating basal cell representations from salient representations associated with perturbation states. This allows for single-cell level generation and *in silico* prediction of perturbation responses. Through evaluations on various datasets, scLAMBDA demonstrated superior performance over existing methods in predicting the effects of both single-gene and two-gene perturbations. Compared to other deep learning-based methods, scLAMBDA offers greater computational efficiency and scalability, making it particularly suitable for handling large-scale datasets. Moreover, scLAMBDA facilitates the identification of upregulated or downregulated genes for specific perturbations and supports the exploration of genetic interaction mechanisms. The scLAMBDA model is available as an open-source Python package at https://github.com/gefeiwang/scLAMBDA, serving as an effective, versatile and efficient resource for single-cell perturbation analysis.

## Results

### Method overview

scLAMBDA models gene expression data from single cells under perturbation conditions in a deep generative learning framework [24, 25], aiming to accurately predict single-cell level gene expression responses to novel perturbations (Fig. 1**a**). This predictive capability is essential for simulating and understanding cellular responses to new experimental conditions.

**Figure 1.**
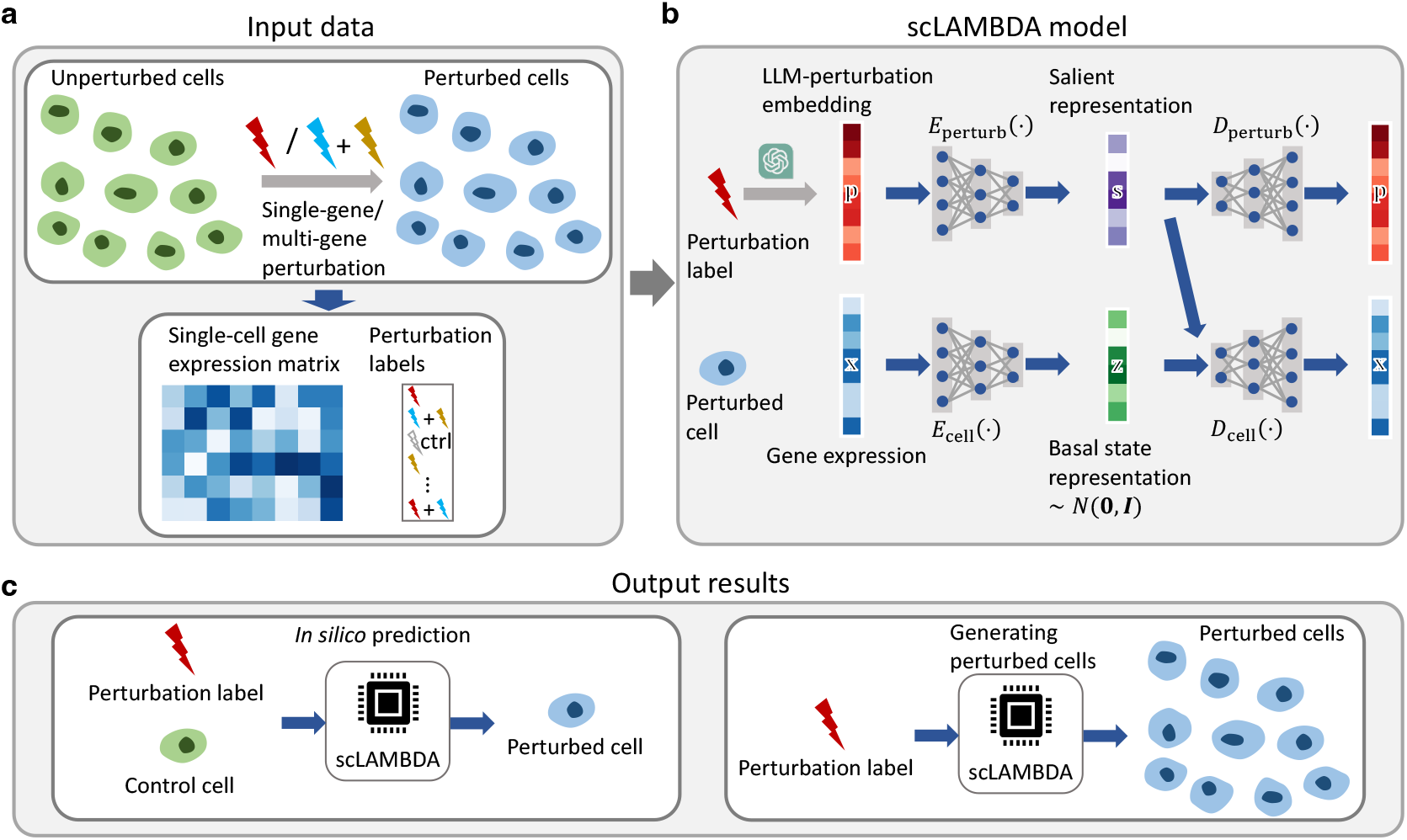
Overview of scLAMBDA. **a**.scLAMBDA takes single-cell genetic perturbation datasets, including the gene expression matrix and perturbation labels, as input. **b**. scLAMBDA employs a deep latent variable framework to model and predict single-cell genetic perturbation responses. It connects genetic perturbation outcomes to embeddings derived from large language models or foundation models and disentangles cell variations into two components: a salient representation encoding perturbation information and a basal cell representation. **c**. Once trained, scLAMBDA enables *in silico* perturbation of individual cells or the generation of cell groups based on specified perturbation information.

To uncover the biological mechanisms underlying perturbation responses, scLAMBDA integrates unsupervised deep representation learning with supervised learning to predict transcriptional outcomes based on high-dimensional perturbation embeddings in its model design. In this framework, each single-cell gene expression profile, represented as a vector **x** ∈ ℝ^*g*^, is associated with a perturbation label containing target genes. Perturbation states are further represented by an embedding vector **p** ∈ ℝ^*p*^, which can be derived from pretrained large language models or foundation models based on target gene names. By default, the embedding vectors obtained from GenePT [22] are utilized in scLAMBDA.

The model applies a deep latent variable approach based on variational autoencoders (VAEs) [26], where gene expression data are generated conditional on both a basal cell state and the perturbation (Fig. 1**b**). Specifically, in the scLAMBDA model, a low-dimensional latent variable with the prior standard normal distribution **z** ∼ 𝒩 (**0, I***d*) captures the basal cell state. In addition to **z**, the model introduces a second embedding, **s** = *E*perturb(**p**) ∈ ℝ^*d*^, referred to as the salient representation, is introduced to encode perturbation-specific information. These two components are combined and decoded through a decoder network to reconstruct gene expression, reflecting the joint contributions of basal cell variability and perturbation effects to changes in gene expression.

To enable accurate post-perturbation gene expression prediction, scLAMBDA introduces disentanglement between basal and perturbation components, as well as adversarial training on perturbation embeddings. Disentanglement is achieved by minimizing mutual information between **z** and **s** using the mutual information neural estimation (MINE) approach [27], allowing distinct representation of basal and salient information. To further enhance model generalization, adversarial examples of **p** are introduced to prevent overfitting in the sparse, high-dimensional perturbation space. This adversarial strategy improves scLAMBDA’s robustness, enabling it to generalize well to unseen perturbations or perturbation combinations by effectively exploring the perturbation space [28].

Finally, scLAMBDA’s training process optimizes the evidence lower bound (ELBO) and mutual information objectives, iteratively refining the generative and disentanglement structures to produce reliable gene expression predictions under varied perturbation conditions. Compared to other deep learning-based methods, such as GEARS, which leverages graph learning, and scGPT, which employs transformers, scLAMBDA offers a more efficient and scalable training process. This is due to its efficient training objective and lightweight network design (Supplementary Tables 1-3), making it well-suited for large-scale datasets.

After training, scLAMBDA enables realistic *in silico perturbation* simulations and the generation of population-level cells based on specific perturbations (Fig. 1**c**), supporting various downstream tasks and assisting in the design of single-cell genetic perturbation experiments. Further details are provided in the Methods section.

### scLAMBDA improved accuracy in predicting single-gene perturbation responses

To evaluate scLAMBDA’s ability to predict single-cell genetic perturbations for unseen single target genes, we first utilized the CRISPR interference (CRISPRi) Perturb-seq dataset from Adamson et al. [29], which targets 86 distinct genes in the human erythroleukemic K562 cell line. We compared scLAMBDA with other state-of-the-art methods for predicting transcriptional responses to genetic perturbations, including GEARS [16], scGPT [18] and GenePert [17], in the experiment.

Using ten random dataset splits (details in the Methods section), we measured the similarity between predicted and observed average changes in gene expression for test perturbations, using the Pearson correlation coefficient (PCC) across top 5,000 highly variable genes. As shown in Fig. 2**a**, scLAMBDA outperformed the other three methods, achieving an average PCC of 0.786 over ten replicates. GenePert followed with an average PCC of 0.775, outperforming GEARS (0.692) and scGPT (0.661). This highlights the strength of linear methods in predicting average gene expression changes. However, PCC evaluates only linear correlations between predicted and observed average gene expression changes, neglecting the heterogeneity of perturbation effects at the cellular level. To assess how well the predicted distributions of perturbed cells resembled the ground truth, we computed the 2-Wasserstein distance (*W*2) [30] as a cell population-level similarity measurement between the predicted and true distributions for each perturbation (details in the Methods section). Notably, GEARS and GenePert are designed primarily for predicting average gene expression changes. For GenePert, we followed GEARS’ approach by adding the predicted average expression changes to a subset of control cells to generate the predicted distribution. As shown in Fig. 2**a**, scLAMBDA produced perturbed cell distributions with the lowest average *W*2 value of 22.552, indicating superior performance in capturing perturbed cell distributions compared to the other methods. We further compared PCC and *W*2 metrics using the top 20 differentially expressed genes based on the *t*-test with Scanpy’s pipeline [31] for each perturbation (Supplementary Fig. 1). All methods achieved comparably high PCC values around 0.8 on differentially expressed genes, but scLAMBDA maintained the lowest *W*2 value of 2.288, showing its effectiveness in accurately modeling perturbed cell distributions.

**Figure 2.**
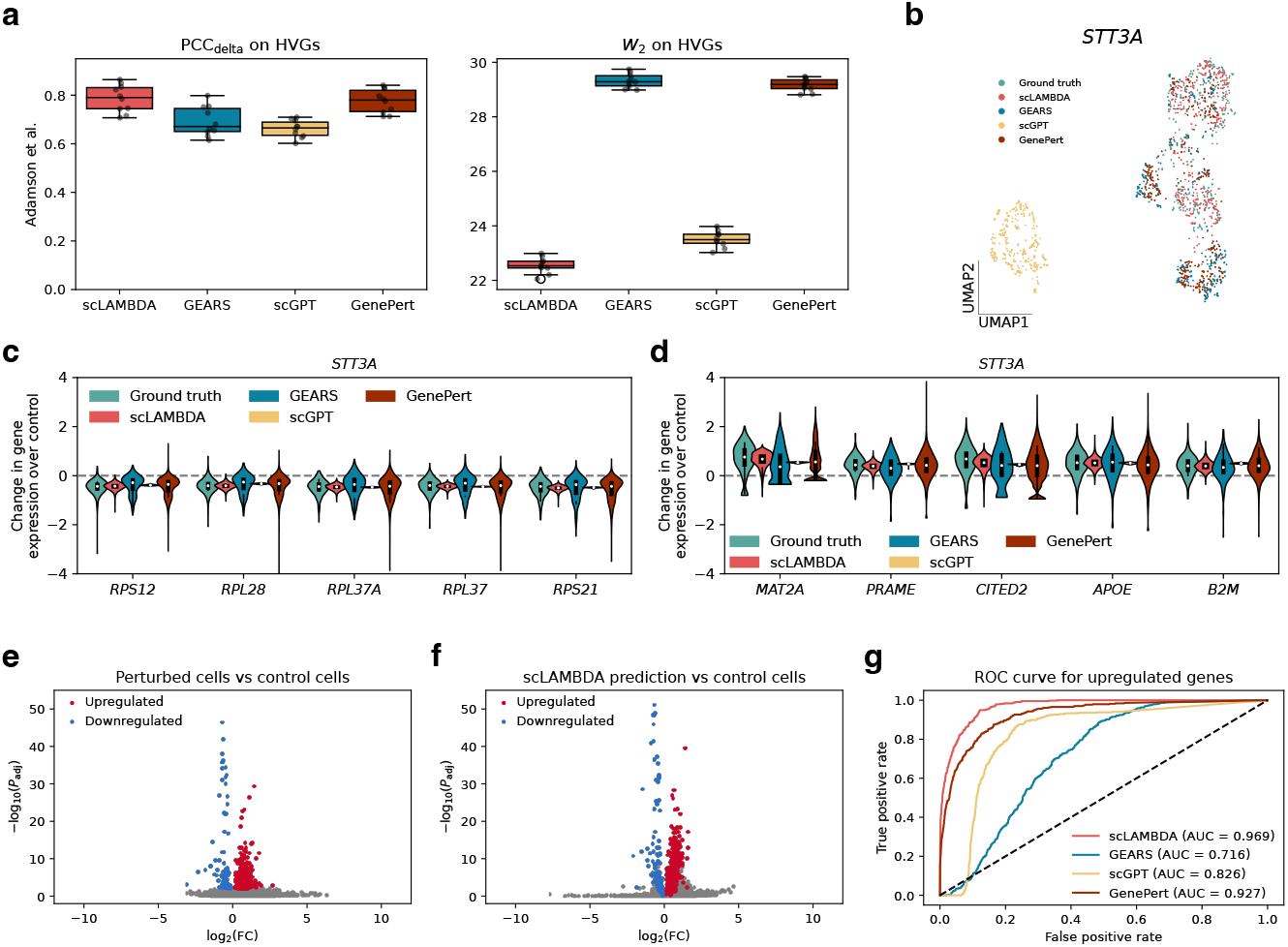
Comparison of predictions for single-gene perturbation outcomes using the Adamson et al. dataset [29]. **a.** Pearson correlation coefficient (PCC) between predicted and true average changes in gene expression (left) and 2-Wasserstein distance (*W*2) between predicted and true distributions (right), evaluated using highly variable genes for ten random dataset splits. HVG, highly variable gene. **b**. UMAP visualization of true and generated cell populations for the knockdown of *STT3A*. **c, d**. Violin plots showing true and predicted post-perturbation gene expression levels for top differentially expressed ribosomal protein genes (**c**) and other genes (**d**) for the knockdown of *STT3A*. **e**. Volcano plot from differential expression analysis comparing true *STT3A* knockdown cells and control cells, highlighting upregulated (log2(FC) *>* 0, adjusted *p*-value *<* 0.01) and downregulated (log2(FC) *<* 0, adjusted *p*-value *<* 0.01) genes in red and blue, respectively. FC, fold change. **f**. Volcano plot from differential expression analysis comparing *STT3A* knockdown cells generated by scLAMBDA and control cells, with upregulated and downregulated genes defined as in **e. g**. Receiver operating characteristic (ROC) curves for identifying upregulated genes as defined in **e** using all methods.

In addition to overall evaluation, we examined the prediction details for individual perturbations. As an example, we visualized the test perturbation targeting the gene *STT3A* using UMAP [32]. In the UMAP plot (Fig. 2**b**), scLAMBDA has the closest alignment to the true perturbation distribution. In comparison, GEARS and GenePert produced results with only partial overlap, while scGPT generated a cell cluster that was distinctly separated from the true perturbation distribution. Since UMAP preserves both local and global structures in single-cell data [33], the well-aligned UMAP clusters further illustrates scLAMBDA’s better prediction quality. Additionally, scLAMBDA accurately predicted the distributions of top differentially expressed genes. For example, gene *STT3A* encodes the catalytic subunit of a complex that interacts with the ribosome [34]. In the Perturb-seq dataset, the knockdown of *STT3A* resulted in the downregulation of several ribosomal protein genes (Fig. 2**c**) and the upregulation of some other genes (Fig. 2**d**). scLAMBDA’s predictions closely captured these true post-perturbation expression levels. By comparison, scGPT tended to underestimate the dispersion of these genes, while GEARS and GenePert failed to recover the distribution shapes of genes such as *MAT2A* and *CITED2*, as their predicted variations primarily reflected the control group. Using scLAMBDA’s generated post-perturbation cells, we are also able to perform differential expression analysis without measuring real cells (details in the Methods section). For *STT3A* knockdown, many upregulated and downregulated genes identified from measured perturbed cells (Fig. 2**e**) served as ground truth. Using scLAMBDA’s results, these genes were reliably detected with a similar differential expression pattern (Fig. 2**f** and Supplementary Fig. 2). Notably, scLAMBDA achieved the highest area under the curve (AUC) scores among all methods, with values of 0.969 for upregulated genes (Fig. 2**g**) and 0.996 for downregulated genes (Supplementary Fig. 3).

### Benchmarking prediction using genome-scale CRISPRi Perturb-seq data

To further demonstrate scLAMBDA’s effectiveness in predicting transcriptional responses to genetic perturbations involving novel target genes, we evaluated all the methods using a genome-scale CRISPRi Perturb-seq dataset from Replogle et al. [35]. This dataset includes 2,393 single essential target genes in cells from the retinal pigment epithelial (RPE1) cell line. Based on numerical metrics, scLAMBDA outperformed all other methods, achieving the highest average Pearson correlation coefficient (PCC = 0.564), surpassing the second best method, GenePert (PCC = 0.547). It also attained the lowest average Wasserstein distance (*W*2 = 28.737), outperforming scGPT (*W*2 = 29.487), on the 5,000 most highly variable genes (Fig. 3**a**). While scGPT ranked second in *W*2 performance on highly variable genes, scLAMBDA consistently delivered lower *W*2 scores across most target genes, as evidenced by results from all ten experimental replicates (Fig. 3**b**). Additionally, scLAMBDA achieved the highest average PCC=0.682 and lowest average *W*2 = 2.715 on the top 20 differentially expressed genes, outperforming all other methods (Supplementary Fig. 4).

**Figure 3.**
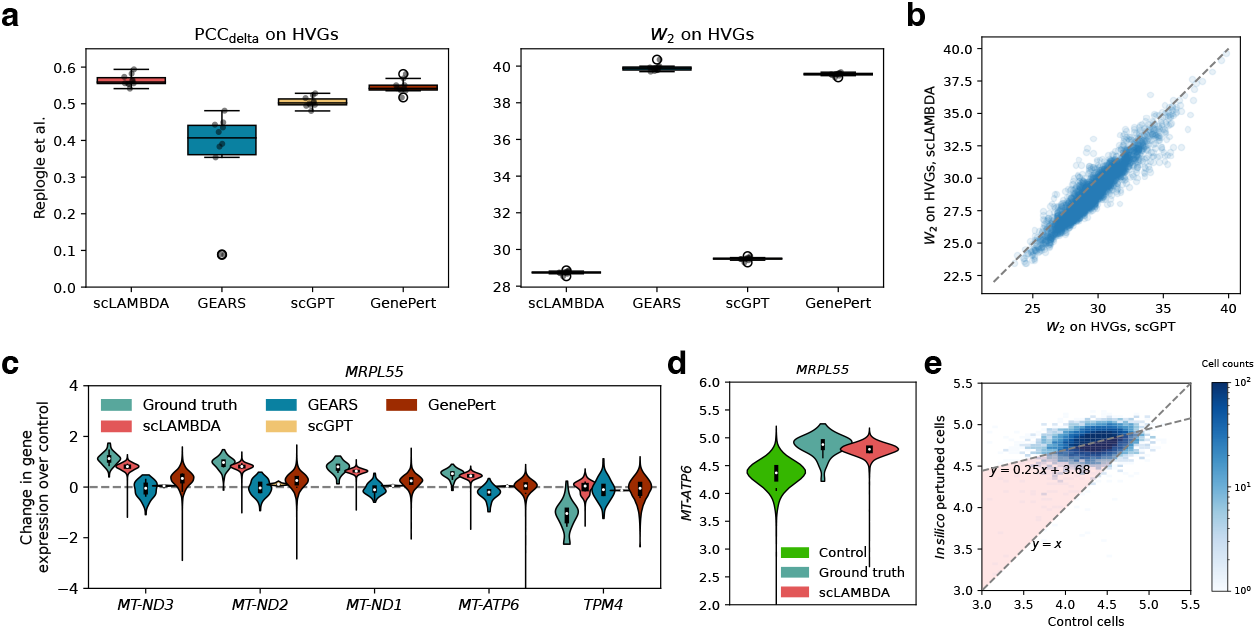
Comparison of predictions for single-gene perturbation outcomes using the Replogle et al. dataset [35]. **a.** PCC between predicted and true average changes in gene expression (left) and *W*2 between predicted and true distributions (right), evaluated using highly variable genes for ten random dataset splits. **b**. Comparison between *W*2 metrics of scLAMBDA and scGPT for all test perturbations across ten replications. **c**. Violin plots showing true and predicted post-perturbation gene expression levels for five top differentially expressed genes for *MRPL55* knockdown. **d**. Violin plots of *MT-ATP6* expression level in control and *MRPL55* knockdown cells. **e**. Density plot comparing the *MT-ATP6* expression levels in control cells with scLAMBDA’s *in silico* predictions for *MRPL55* knockdown.

Using this dataset, we also illustrated the interpretability of scLAMBDA’s *in silico* perturbation result. For example, one focus of the original study was to investigate the transcriptional responses of the mitochondrial genome to different mitochondrial stressors [35]. Among these stressors, knocking down the test target gene *MRPL55*, which encodes a 39S subunit protein in mitochondrial ribosomes, was found to upregulate several mitochondrial genes, as evidenced by the expression profile of perturbed cells (Fig. 3**c**). Unlike other methods, scLAMBDA accurately predicted the perturbed distributions of these genes. For instance, gene *MT-ATP6* exhibited a higher mean and lower dispersion compared to the control expression in its perturbed expression compared to the control in the original dataset (Fig. 3**d**). However, without paired measurements, the perturbation effect on individual cells is challenging to discern. Using the orthogonal distance regression which considers errors in both the measurement of explanatory variables and responses [36], we fitted the relationship between the control group expressions and scLAMBDA’s predicted expressions for control cells. scLAMBDA’s predictions revealed that the upregulation of *MT-ATP6* was heterogeneous across cells, with smaller increases observed at higher original expression levels, eventually saturating beyond a specific threshold (Fig. 3**e**). The above results further highlighted scLAMBDA’s reliability and utility in predicting single-cell perturbation responses for novel single target genes.

### Prediction of multi-gene perturbation effects

After validating scLAMBDA’s predictions for novel single-gene perturbation effects, we next assessed its ability to predict multi-gene perturbation effects using the CRISPR activation (CRISPRa) Perturb-seq dataset of K562 cells from Norman et al. [37], which includes 104 single-gene perturbations and 130 two-gene perturbations. The combinatorial effects of multiple target genes are often non-additive, reflecting complex genetic interactions that are critical for understanding cellular function [37, 38]. These interactions reveal how genes influence each other to regulate cellular processes, improving our understanding of the properties that arise when genes are expressed together. Predicting perturbation effects in multi-gene scenarios requires methods that can not only generalize to unseen target genes but also effectively capture the combinatorial interactions between different target genes.

As illustrated in the UMAP plot of post-perturbation gene expressions, the perturbations induced distinct shifts relative to the control cell group (Fig. 4**a, b**). After training, we visualized the representations learned by scLAMBDA. The UMAP plot in Fig. 4**c, d** shows that scLAMBDA effectively integrates cells with different perturbation labels into its learned basal cell representation.

**Figure 4.**
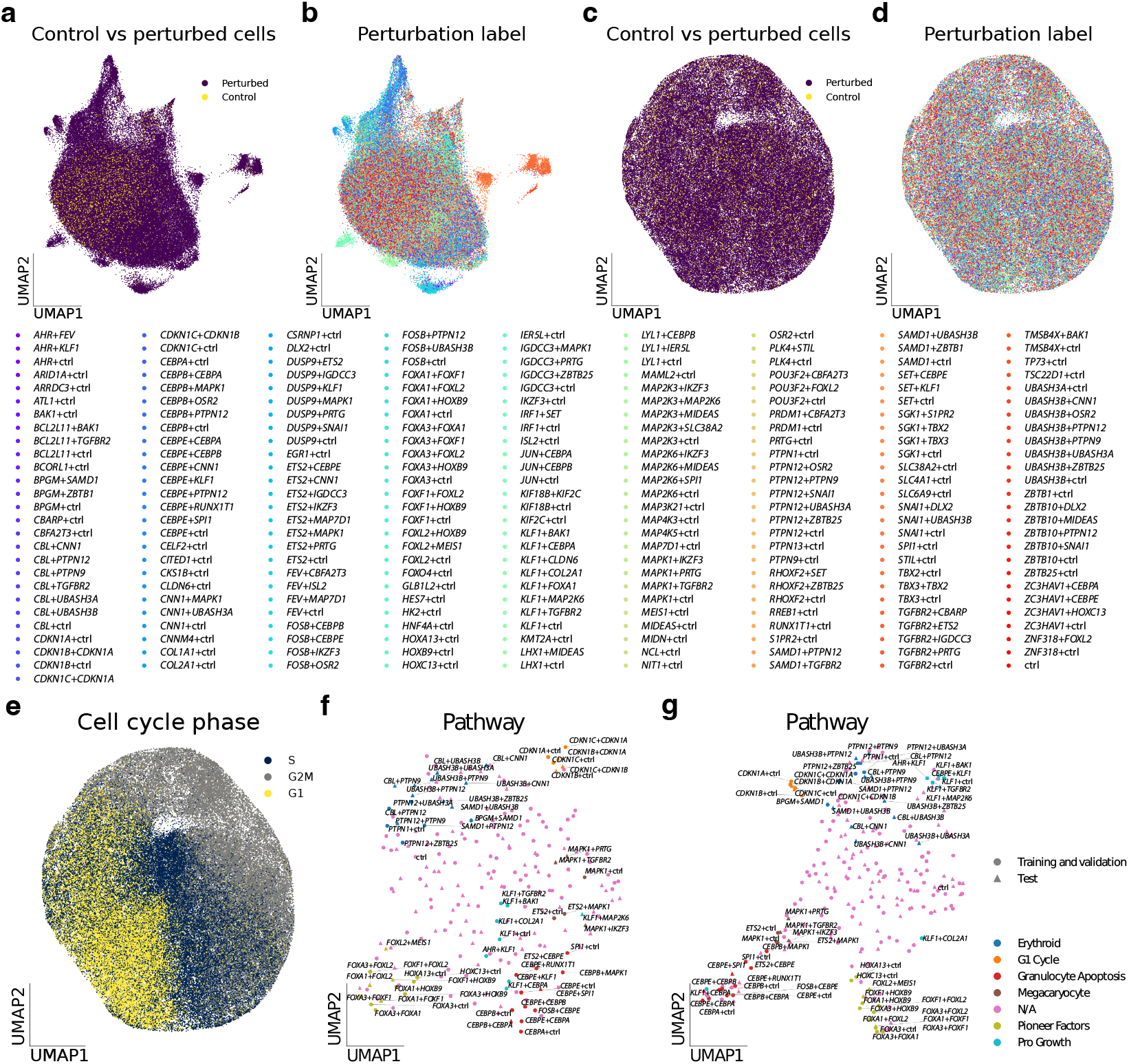
Disentangled representation learning with scLAMBDA. **a-b.** UMAP plots of the CRISPRa Perturb-seq dataset from Norman et al. [37], colored by control vs. perturbed status (**a**) and target gene labels (**b**). **c**-**e**. UMAP plots of scLAMBDA’s learned basal representations, colored by control vs. perturbed status (**c**), target gene labels (**d**), and cell cycle phases (**e**). **f**. UMAP plot of perturbation embeddings generated by GenePT, colored by perturbation clusters. **g**. UMAP plot of salient representations learned by scLAMBDA, colored by perturbation clusters.

To further explore the variation captured by the basal cell representation, we calculated cell cycle gene scores and assigned cell cycle phases to all cells [39, 31]. The results, shown in Fig. 4**e**, reveal that the basal cell representation encapsulates cell cycle information, underscoring scLAMBDA’s capability to learn fundamental variations underlying cellular states.

Next, we examined the salient representation learned by scLAMBDA. We visualized UMAP plots of perturbation embeddings generated by GenePT [22] (Fig. 4**f**), which serve as the default input to scLAMBDA, and the salient representation produced by scLAMBDA (Fig. 4**g**). The salient representation, directly encoded from the perturbation embeddings and refined during training to reconstruct post-perturbation gene expression, acts as an intermediate variable linking large language model-based prior knowledge with transcriptional responses to perturbations. The UMAP plots were colored based on perturbation clusters identified by Norman et al. [37], using the similarity of post-perturbation gene expression profiles. As shown in Fig. 4**g**, the salient representation generalized effectively to test perturbations by incorporating large language model-derived prior information. Additionally, compared to the UMAP of the initial perturbation embeddings, the salient representation integrated gene expression data and displayed more compact clustering of transcriptionally similar perturbations, highlighting its improved alignment with biological responses.

To comprehensively evaluate the performance of all methods, we divided the test perturbations into four categories, including 0/1 seen, 0/2 seen, 1/2 seen, and 2/2 seen, based on their overlap with the training set (Fig. 5**a**). We evaluated the overall prediction performance of all methods using the PCC and *W*2 metrics and further analyzed their performance across each category. For this dataset, scLAMBDA demonstrated superior overall prediction accuracy, achieving the highest average PCC=0.664 and lowest average *W*2 = 13.228 on highly variable genes. GenePert followed with an average PCC of 0.635, while scGPT achieved the second lowest *W*2 of 13.912 (Fig. 5**b**). Besides, scLAMBDA also showed the best overall accuracy evaluated on differentially expressed genes (Supplementary Fig. 5). For each individual category, scLAMBDA outperformed other methods in both PCC and *W*2 metrics on highly variable genes (Fig. 5**c**-**f**). In the three scenarios of two-gene combination, scLAMBDA demonstrated progressively higher average PCCs 0.593, 0.718, and 0.850, as the number of target genes seen in the training set increased from zero to one and then to two. These results suggest that while scLAMBDA generalizes well to unseen target genes, the prediction accuracy for multi-gene combinatorial perturbation effects could be enhanced if the effects of more individual target genes in the combination are known.

**Figure 5.**
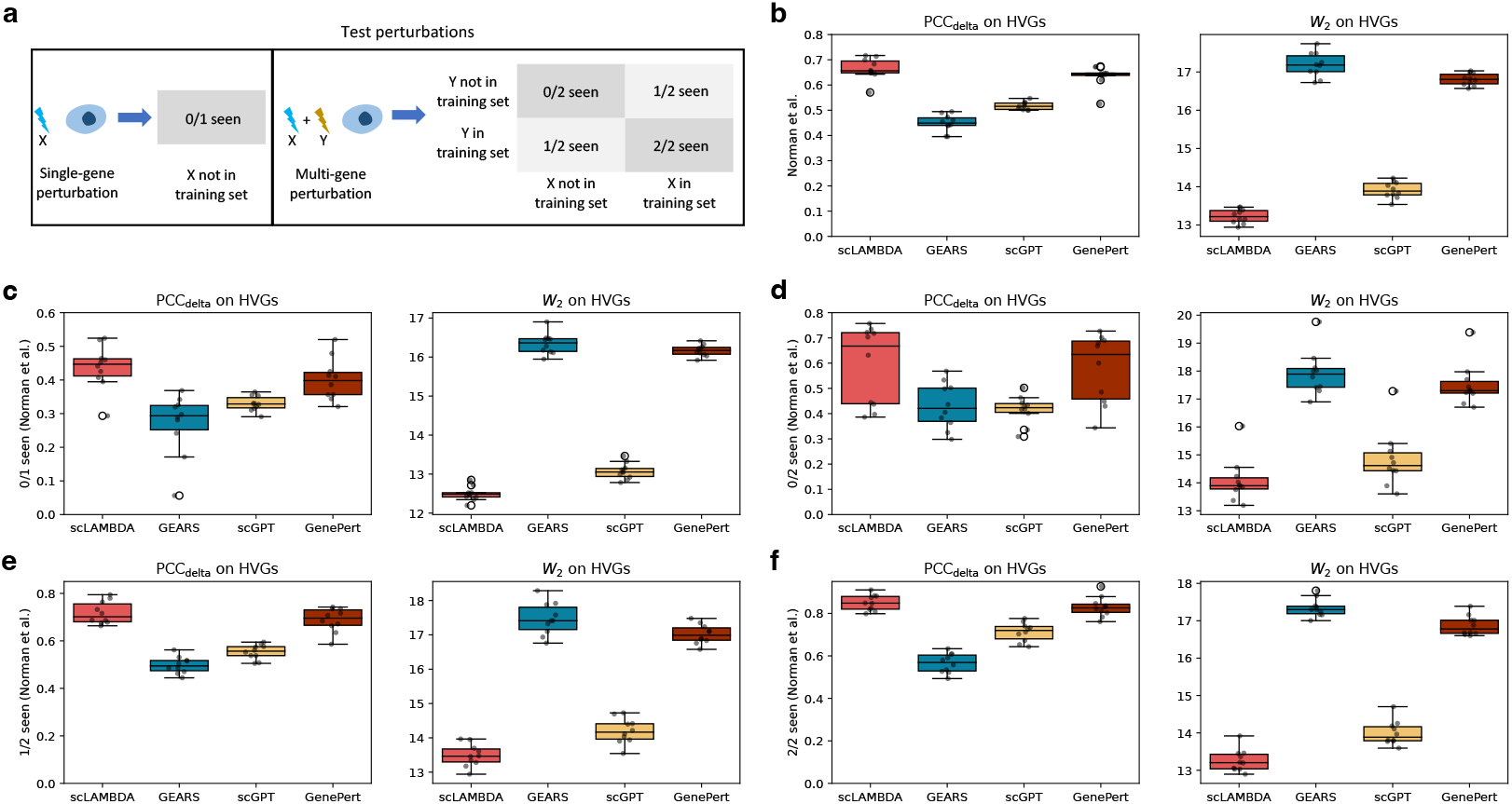
Comparison of methods for predicting multi-gene perturbation outcomes. **a.** Test perturbation categories. **b**-**f**. Performance comparison across all test perturbations (**b**), single-gene test perturbations (**c**), and two-gene test perturbations with zero (**d**), one (**e**), or two (**f**) genes observed during training. Methods are evaluated based on PCC between predicted and true average changes in gene expression (left) and *W*2 between the predicted and true distributions (right), using highly variable genes for ten random dataset splits.

Using this dataset, we further validated the contributions of scLAMBDA’s core model components. The analysis revealed that the inclusion of mutual information regularization and adversarial training effectively improved prediction accuracy (Supplementary Fig. 6). Additionally, we assessed the model’s performance using different gene embeddings, including GenePT, scGPT, and two genome sequence foundation models, HyenaDNA [40] and DNABERT-2 [41]. For a fair comparison, we utilized the gene embeddings from scGPT’s pretrained model without fine-tuning on the Perturb-seq dataset. As illustrated in Supplementary Fig. 7, scLAMBDA achieved the second highest performance when using scGPT gene embeddings, surpassed only by the results obtained with GenePT gene embeddings. In contrast, embeddings derived from genome sequence foundation models demonstrated suboptimal accuracy, indicating that the genome sequence similarity between target genes is less informative for the prediction of genetic perturbation effects.

### scLAMBDA facilitates the exploration of genetic interactions

Building on its high accuracy in predicting multi-gene perturbation effects, we next explored scLAMBDA’s capability to map the landscape of genetic interactions using its predictions for two-gene combinations. To demonstrate scLAMBDA’s utility, we evaluated its accuracy across key genetic interaction scores such as magnitude, model fit, and singles to double similarity, which are critical for defining genetic interaction types [37] (details in the Methods section). For instance, two genes are classified as synergistic when their combined magnitude exceeds one, and as suppressive when the magnitude is low. To assess the precision of genetic interaction predictions across all methods, we calculated the root mean square error (RMSE) between predictions and ground truth for two-gene perturbations, focusing on cases where both target genes were included in the training data for each experimental replicate. As shown in Fig. 6**a**, scLAMBDA achieved the lowest mean RMSE for these scores, highlighting its robustness and reliability in predicting genetic interactions.

**Figure 6.**
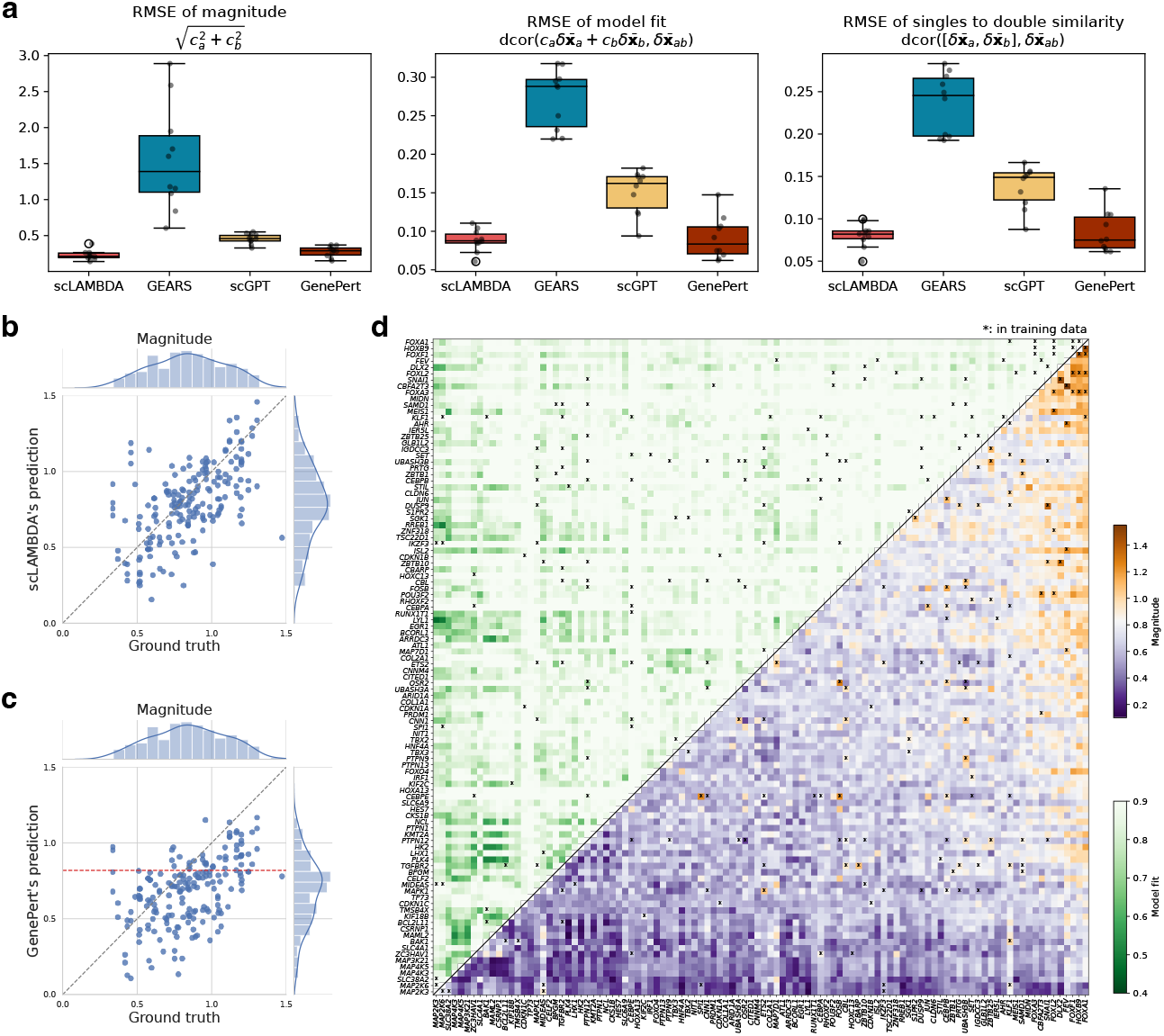
Genetic interaction prediction. **a.** Prediction accuracy of genetic interaction scores including magnitude (left), model fit (middle) and singles to double similarity, measures using root mean square error (RMSE). **b**. Comparison between scLAMBDA’s prediction of magnitude with ground truth across ten replications. **c**. Comparison of GenePert’s predicted magnitudes with ground truth across ten replicates. The red dashed line represents the theoretical value of 0.819, around which GenePert’s predictions are expected to concentrate. **d**. Genetic interaction map predicted by scLAMBDA, showing magnitude and model fit for all two-gene combinations.

Although GenePert achieved the second highest genetic interaction score prediction accuracy based on RMSE, as a linear model, it may introduce bias when estimating genetic interaction scores. This is because GenePert assumes that the average post-perturbation gene expression is a linear function *fθ* of the perturbation embeddings, such that for a single target gene *a*, 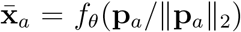. For perturbations involving two target genes, *a* and *b*, this assumption extends to 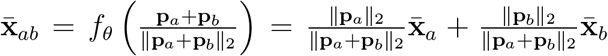 Therefore, the mean change in gene expression follows 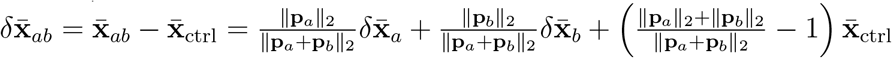, where 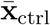 is the mean of gene expressions of control cells. GenePert also uses GenePT gene embeddings, where the 2-norm is one for individual genes and concentrates around a mean of 1.726 for two-gene combinations (Supplementary Fig. 8). If the perturbation effects are orthogonal to the mean control cell gene expression, through straightforward mathematical derivations, the coefficients estimated from GenePert’s results would tend to be similar and close to 0.579, with the estimated magnitudes biased toward approximately 0.819. This pattern was indeed observed in practice, as reflected in the coefficients and magnitudes derived from GenePert’s predictions (Fig. 6**b, c** and Supplementary Fig. 9). Unlike GenePert, scLAMBDA avoids this issue by employing nonlinear neural networks to model the relationship between perturbation embeddings and gene expression.

Building on the demonstrated accuracy of scLAMBDA’s genetic interaction predictions, we trained the model on all 104 single-gene and 130 two-gene perturbations to predict genetic interactions for all possible gene pairs. scLAMBDA extends the genetic interaction map to all two-gene combinations by leveraging gene similarity embedded in the prior information from gene embeddings (Fig. 6**d**). For instance, the training data showed synergistic combinatorial effects between *POU3F2* and *FOXL2*. Using similarity among genes encoding FOX proteins, scLAMBDA also predicted potential synergistic effects between *POU3F2* and other FOX family members, such as *FOXA1* and *FOXF1*, highlighting its ability to capture biologically meaningful relationships for genetic interaction prediction.

## Discussion

In this paper, we introduced scLAMBDA, a deep generative learning framework designed to model and predict single-cell transcriptional responses to genetic perturbations, including both single-gene and combinatorial multi-gene scenarios. scLAMBDA demonstrated robust performance across multiple datasets, consistently outperforming state-of-the-art methods such as GEARS, scGPT, and GenePert. Its ability to generalize to unseen genes and perturbations with high predictive accuracy, underscores its value in guiding experimental designs for genetic perturbation studies.

By leveraging the embedding capabilities of large language models and employing a disentangled representation learning framework, scLAMBDA addresses key limitations in existing computational methods. Incorporating gene embeddings derived from pretrained language models, scLAMBDA effectively integrates prior biological knowledge into its predictions. Additionally, the accessibility of gene embeddings from large language models and foundation models [22, 42] ensures that scLAMBDA is broadly applicable to diverse target genes. In contrast, GEARS relies on the presence of target genes within a gene ontology-based knowledge graph, and scGPT requires target genes to have measured expression data in sequencing datasets, limiting their applicability. Additionally, unlike methods that focus primarily on average gene expression changes such as GEARS and GenePert, scLAMBDA utilizes the disentangled generative learning framework to model the heterogeneity of single-cell responses, providing more insights into cellular behavior and enhances the interpretability of perturbation effects. Furthermore, scLAMBDA offers superior computational efficiency during training compared to other deep learning-based methods (Supplementary Fig. 10), benefiting from its simple input format and flexible model architecture.

While scLAMBDA represents a significant advancement in single-cell perturbation modeling, there remain several limitations. For example, the model’s performance is influenced by the quality of gene embeddings (Supplementary Fig. 7), suggesting that further refinement and exploration of embedding strategies could enhance its predictive accuracy. Additionally, scLAMBDA’s framework has the potential to be extended to other types of perturbations with effective embeddings, such as chemical perturbations, which is a promising direction for future exploration.

In conclusion, scLAMBDA shows a significant advancement for predictive modeling in single-cell genetic perturbation analysis. Its ability to integrate prior knowledge, capture single-cell heterogeneity, and generalize to novel perturbations makes it a versatile tool for functional genomics research. As computational models continue to play an increasingly central role in biology, we believe scLAMBDA’s framework will provide valuable insights for advancing single-cell perturbation studies.

## Methods

### The model of scLAMBDA

Let **x** ∈ ℝ^*g*^ represent the gene expression vector of a single cell, where the associated labels specify the target gene of the perturbation. For each perturbation label, we define an embedding **p** ∈ ℝ^*p*^, which can be obtained from a pretrained model using the target gene name. For perturbations involving two target genes *a* and *b*, the gene embedding is calculated as the sum of the individual gene embeddings: **p***ab* = **p***a* + **p***b*.

### Latent variable modeling

In the scLAMBDA framework, we model the gene expression of a single cell using the following generative model:

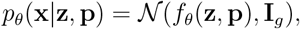

where **z** ∼ 𝒩 (**0, I***d*) is a low-dimensional latent variable representing the basal cell state. The mean function *fθ*(**z, p**) = *D*cell(**z** + **s**) is parameterized such that **s** = *E*perturb(**p**), where *E*perturb(·) is a neural network that encodes the high-dimensional perturbation embedding **p** into a lower-dimensional salient representation vector **s**, with the same dimensionality as **z**. The network *D*cell(·) then decodes **z** and **s** into the mean of the gene expression **x**.

Since the marginal distribution *pθ*(**x**|**p**) = ∫ *pθ*(**x**|**z, p**)*p*(**z**) d**z** is intractable, we approximate it using a variational autoencoder (VAE) approach [26]. Specifically, we parameterize a variational posterior distribution: *qϕ*(**z**|**x**) ≈ *pθ*(**z**|**x**) = *pθ*(**z**|**x, p**), where the equality holds because we assume that the basal cell state **z** does not involve perturbation information. The variational posterior is modeled as a Gaussian distribution *qϕ*(**z**|**x**) = 𝒩 (***µ***, diag(***σ***2)), where the mean ***µ*** and variance ***σ*** are encoded from **x** as (***µ***, log ***σ***) = *E*cell(**x**) via an encoder network *E*cell(·).

We can then train the model by optimizing the evidence lower bound (ELBO):

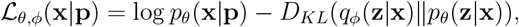

which can be expressed as:

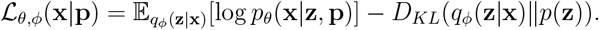

### Disentanglement of the basal and salient representations

In the model described above, when the latent variable **z** is encoded from the input **x**, it may still be dominated by perturbation-related variations, even after optimizing the ELBO. To accurately separate the biological variation in single-cell perturbation data into basal cell state and salient representation, we introduce an additional decoder network, *D*perturb(·). This network ensures that the perturbation embedding **p** is fully captured in the salient representation **s** by minimizing the reconstruction error 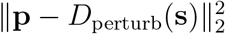.

To further disentangle **z** (basal representation) and **s** (salient representation), we aim to minimize the mutual information (MI) between these two variables, defined as:

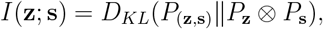

where *P***z** ⊗ *P***s** represents the product of their marginal distributions. Since directly computing *I*(**z**; **s**) is intractable, we adopt the mutual information neural estimation (MINE) approach [27] for its estimation. Specifically, MINE is formulated as:

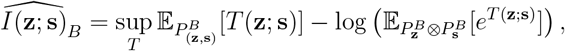

where *T* (**z**; **s**) ∈ ℝ is a scalar function, and *PB* represents an empirical distribution based on *B i*.*i*.*d*. samples from *P*. During training, we parametrize *T* as an auxiliary network *Tψ*(**z**; **s**) and iteratively optimize it along with the other networks to estimate and minimize *I*(**z**; **s**), promoting the disentanglement of the basal and salient representations. When minimizing *I*(**z**; **s**), we only train **z** and keep **s** fixed with the stop gradient operator. Additionally, since **z** is sampled from *qϕ*(**z**|**x**) = 𝒩 (***µ***, diag(***σ***2)), we replace **z** with ***µ*** in MINE, ignoring the additional independent noises.

### Improving generalization of gene expression prediction by adversarial training on perturbation embeddings

To address potential overfitting and improve generalization to unseen perturbations **p**, scLAMBDA introduces adversarial examples into the distribution of **p**. Since scLAMBDA generates post-perturbation gene expression **x** ≈ *D*cell(**z** + *E*perturb(**p**)) based on **p**, there is a risk of overfitting due to the sparse nature of **p** in high-dimensional space. With each dataset containing only a limited number of genetic perturbation types, this sparsity can lead to poor generalization.

To address this issue, we augment the distribution of **p** with adversarial examples [28]. For each cell **x** and its corresponding perturbation **p**, we first generate the reconstructed cell 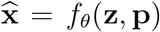 with **z** ∼ *qϕ*(**z**|**x**). We then create an adversarial example 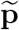, which acts as a “worst-case” perturbation that remains near **p** but maximizes prediction error:

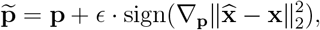

where the fast sign gradient method [43] is applied to **p** to increase the prediction error. We set *ϵ* = 0.001∥**p**∥ by default to ensure 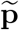 remains close to **p**.

### Training algorithm of scLAMBDA

We summary the training procedure of scLAMBDA in Algorithm 1. We used the Adam optimizer [44] for the gradient descent updates. After training, scLAMBDA can perform *in silico* perturbation experiment for cells or generate a new population of cells with a given perturbation.

#### Algorithm 1

Training algorithm of scLAMBDA.

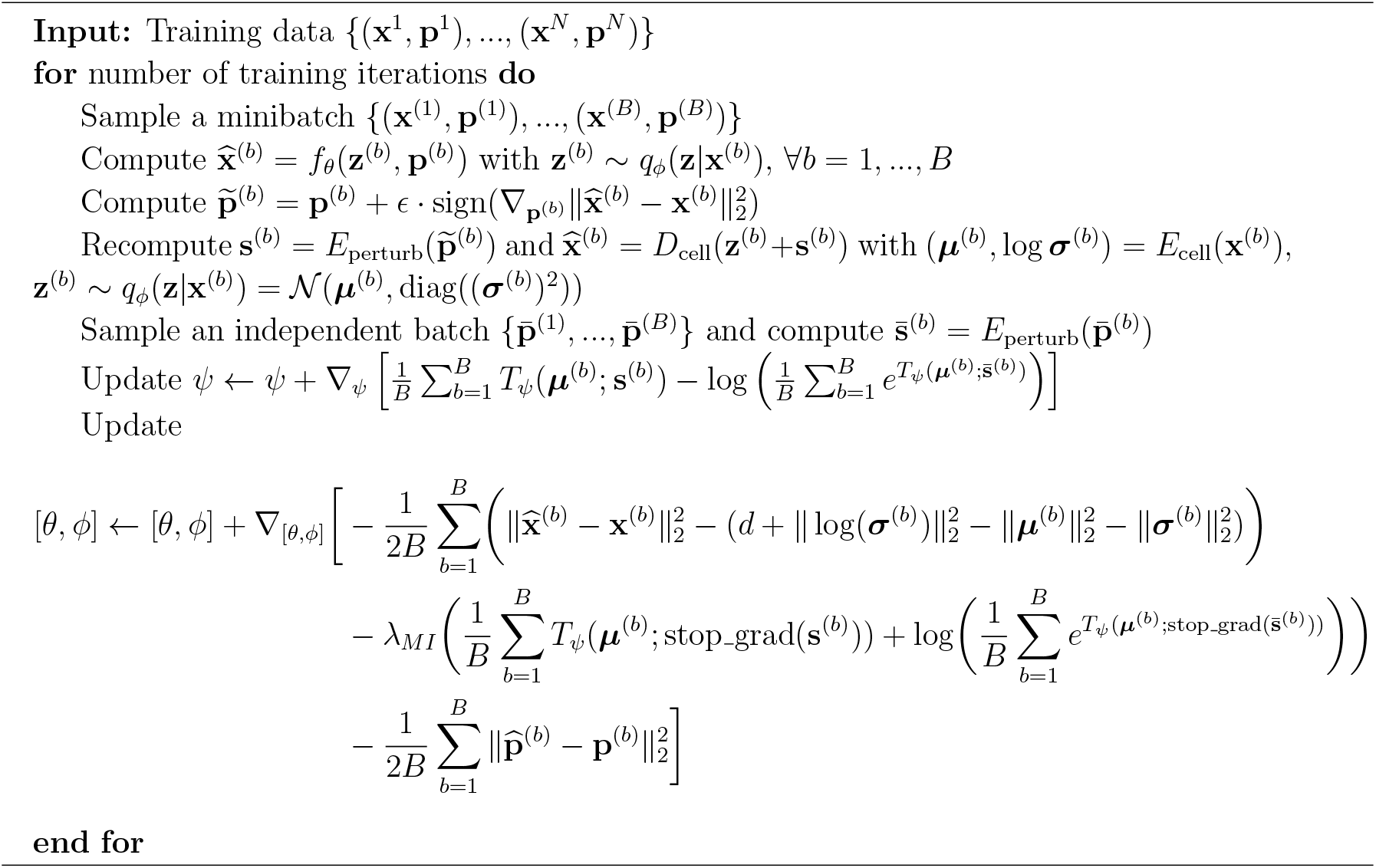

## Analysis details

### Model training details

For all datasets, the model was trained using the top 5,000 highly variable genes in all cells. By default, scLAMBDA employs a batch size of 500. The optimization process spans 200 epochs, starting with an initial learning rate of *lr* = 0.0005, which is scaled by a factor of *γ* = 0.2 every 30 epochs. The coefficients for running averages are set to (*β*1, *β*2) = (0.9, 0.999). The dimensionality of the latent space is set to *d* = 30, and the regularization parameter is set to *λMI* = 200.

### Evaluation metrics

*Pearson correlation coefficient (PCC)*. For each test perturbation, we measure the similarity between the mean true and predicted gene expression changes. Let 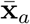 and 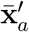 be the means of true and predicted gene expressions for perturbation *a*, and 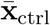 be the mean of gene expressions of control cells. Denote 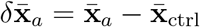 and 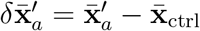. Then the PCC is calculated as

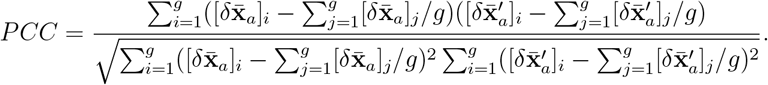

*2-Wasserstein distance (W*2). Given two empirical distributions with samples **x**1, …, **x***n* and **y**1, …, **y***n*, the *W*2 metric computes the optimal cost of transporting one set of samples to the other:

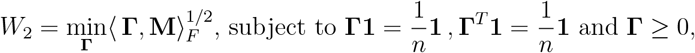

where **M** is the cost matrix defined as 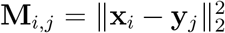. For efficient computation, we sample *n* = 100 cells from true perturbed cells and generated cells for all methods.

### Data splitting

For single-gene perturbation datasets, the data were initially split into 75% training perturbations and 25% test perturbations. Within the training data, 10% of the perturbations were designated as validation data to avoid overfitting based on the highest validation accuracy, measured by PCC. For two-gene perturbation datasets, we followed the GEARS protocol by first dividing all target genes into 75% training genes and 25% test genes. The test set included all single-gene perturbations targeting test genes, all two-gene perturbations involving at least one test gene, and 25% of two-gene perturbations where both target genes were in the training set. The same procedure, using a 90% and 10% split of target genes, was applied to separate training and validation data.

### Differential expression analysis

Since different methods generate varying numbers of cells for each perturbation, we subsampled *n* = 300 cells as the predicted perturbed cell population for each method to eliminate the effect of sample size on statistical inference. We then performed a *t*-test for differential expression analysis between the generated cell populations and the control cells. Genes with adjusted *p*-values *<* 0.01 from the comparison between true perturbed cells and control cells were considered ground truth upregulated or downregulated genes, with fold change (FC) values greater than one for upregulated genes and less than one for downregulated genes.

### Genetic interaction scores

We followed the approach of Norman et al. [37] to define genetic interaction scores for two target genes, *a* and *b*, based on the model 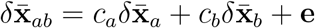, where 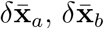, and 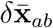 represent the mean gene expression changes induced by perturbation of single target gene *a*, single target gene *b*, and the combination of target genes *a* and *b*, respectively. To compute *ca* and *cb*, we performed a regression of 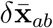 on 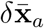 and 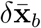, focusing on highly variable genes with a minimum of 100 counts and expression in at least three cells. Once *ca* and *cb* are computed, we calculated genetic interaction scores, including magnitude 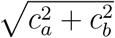 model fit 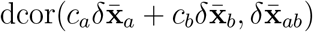 and singles to double similarity 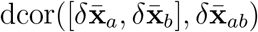, where function dcor(·) represents distance correlation [45]. Specifically, a large magnitude indicates potential synergy between genes, while a small magnitude indicates suppression. Besides, a small model fit indicates neomorphism, and a large singles to double similarity indicates redundancy [37, 16].

### Computational time and memory usage

We evaluated the computational time and memory usage of all methods using the Norman et al. dataset [37]. As shown in Supplementary Fig. 10, scLAMBDA demonstrated superior computational efficiency compared to other deep learning models. Its training time was only longer than GenePert, which is a ridge regression model trained on average post-perturbation gene expression changes.

## Data availability

All data used in this work are publicly available through online sources.

- Perturb-seq dataset from Adamson et al. [29] (GSE90546).
- Perturb-seq dataset from Replogle et al. [35] (GSE146194).
- Perturb-seq dataset from Norman et al. [37] (GSE133344).

## Code availability

scLAMBDA software is available at https://github.com/gefeiwang/scLAMBDA.

## Acknowledgements

This work was supported in part by NIH grants R01 GM134005, U24 HG012108 and P50 CA196530.

