## Supplementary Information for "Modeling and predicting single-cell multi-gene perturbation responses with scLAMBDA"

#### Supplementary Figures

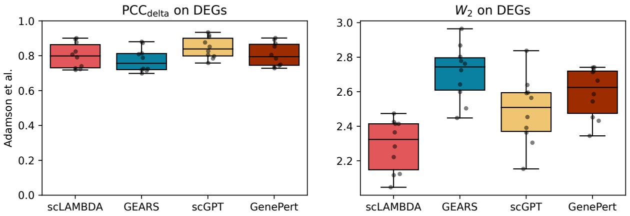


Supplementary Fig. 1: **Comparison of predictions for single-gene perturbation outcomes using the Adamson et al. dataset [1] on top differentially expressed genes (DEGs).** Pearson correlation coefficient (PCC) between predicted and true average changes in gene expression (left) and 2-Wasserstein distance ($W_2$) between predicted and true distributions (right), evaluated using DEGs for ten random dataset splits.

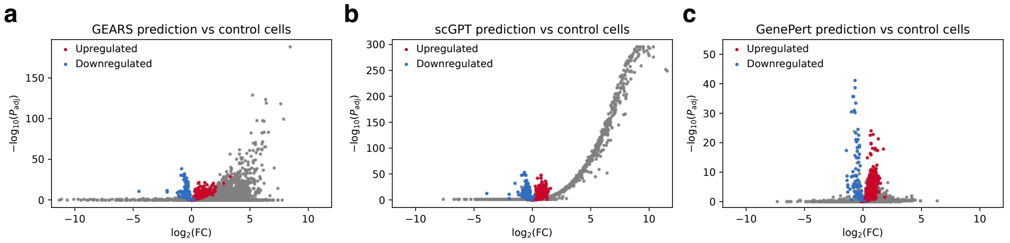


Supplementary Fig. 2: **Volcano plots from differential expression analysis for *STT3A* knockdown. a-c**. Volcano plots from differential expression analysis comparing *STT3A* knockdown cells generated by GEARS (**a**), scGPT (**b**) or GenePert (**c**) and control cells, with upregulated and downregulated genes defined by the differential expression analysis comparing true *STT3A* knockdown cells and control cells.

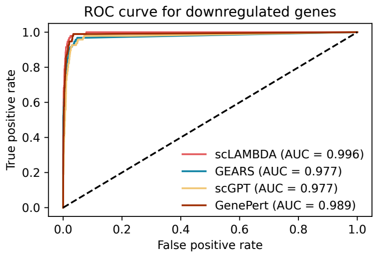


Supplementary Fig. 3: **Receiver operating characteristic (ROC) curves for identifying downregulated genes defined by the differential expression analysis comparing true *STT3A* knockdown cells and control cells.**

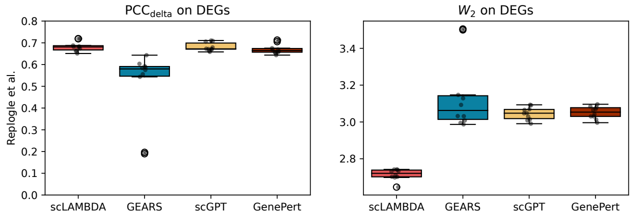


Supplementary Fig. 4: **Comparison of predictions for single-gene perturbation outcomes using the Replogle et al. dataset [2] on top DEGs.** PCC between predicted and true average changes in gene expression (left) and $W_2$ between predicted and true distributions (right), evaluated using DEGs for ten random dataset splits.

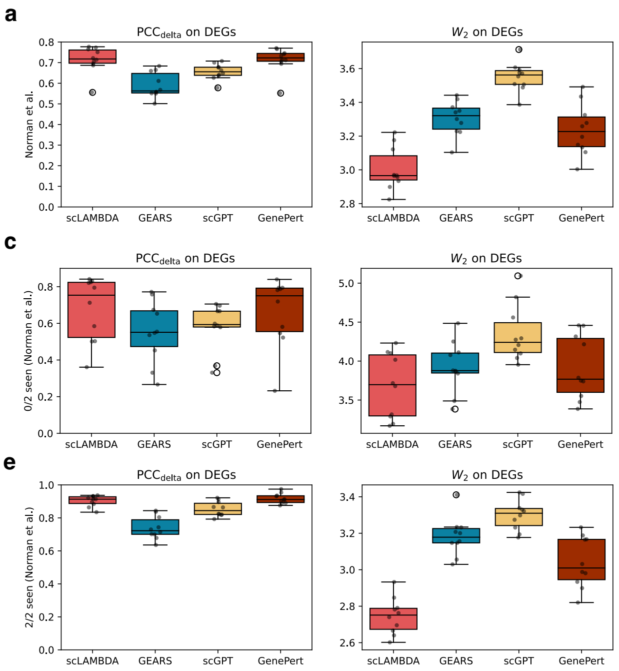


Supplementary Fig. 5: **Comparison of methods for predicting multi-gene perturbation outcomes using the Norman et al. dataset [3] on top DEGs. a-e.** Performance comparison across all test perturbations (**a**), single-gene test perturbations (**b**), and two-gene test perturbations with zero (**c**), one (**d**), or two (**e**) genes observed during training. Methods are evaluated based on PCC between predicted and true average changes in gene expression (left) and $W_2$ between predicted and true distributions (right), using DEGs for ten random dataset splits.

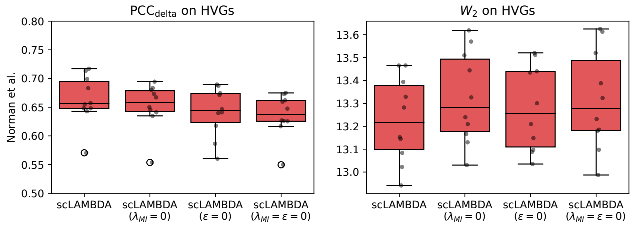


Supplementary Fig. 6: **Evaluation of scLAMBDA's performance by removing its components.** We used the Perturb-seq data from Norman et al. [3] evaluate scLAMBDA's performance by removing its components, based on PCC between predicted and true average changes in gene expression (left) and $W_2$ between predicted and true distributions (right), using HVGs for ten random dataset splits. Notably, scLAMBDA showed less satisfactory performance when removing the mutual information regularization ($\lambda_{MI} = 0$) or removing adversarial training ($\epsilon = 0$).

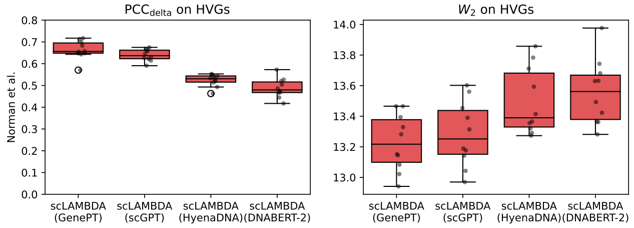


Supplementary Fig. 7: **Evaluation of scLAMBDA's performance using different gene embeddings.** We used the Perturb-seq data from Norman et al. [3] evaluate scLAMBDA's performance using gene embeddings obtained from GenePT [4], scGPT [5], HyenaDNA [6] and DNABERT-2 [7], based on PCC between predicted and true average changes in gene expression (left) and $W_2$ between predicted and true distributions (right), using HVGs for ten random dataset splits.

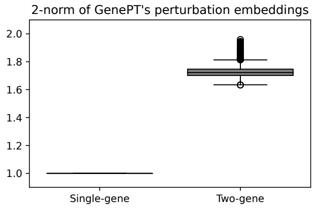


Supplementary Fig. 8: **2-norm distributions of perturbation embeddings.** We calculated the 2-norms of perturbation embeddings of single genes and combinations of two genes, using gene embeddings obtained from GenePT.

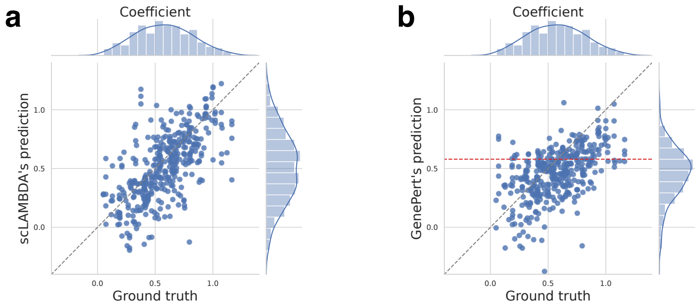


Supplementary Fig. 9: **Comparison between predictions of coefficients with ground truth across ten replications.** **a**. Comparison between scLAMBDA's prediction of coefficients ($c_a$ and $c_b$) with ground truth across ten replications. **b**. Comparison between GenePert's prediction of coefficients with ground truth across ten replications. The red dashed line represents the theoretical value of 0.579, around which GenePert's predictions are expected to concentrate.

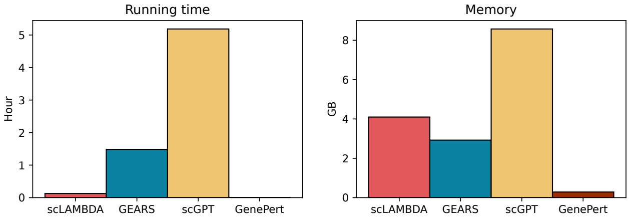


Supplementary Fig. 10: **Computational time and peak memory usage.** We used the Perturb-seq data from Norman et al. [3] to compare the computational time (left) and peak memory usage (right) of all compared methods.

### Supplementary Tables

#### Network structures in scLAMBDA

| Layer | Detail | Input Size | Output Size | Number of Parameters |
|---|---|---|---|---|
| Fully Connected | Linear | $5,000$ | 512 | $512 \times 5,000 + 512 = 2,565,000$ |
| | Leaky ReLU | 512 | 512 | 0 |
| Fully Connected | Linear | 512 | 512 | $512 \times 512 + 512 = 262,656$ |
| | Leaky ReLU | 512 | 512 | 0 |
| Fully Connected | Linear | 512 | $30 \times 2$ | $512 \times 60 + 60 = 30,780$ |

Supplementary Table 1: **Structure of scLAMBDA's encoder networks.**

| Layer | Detail | Input Size | Output Size | Number of Parameters |
|---|---|---|---|---|
| Fully Connected | Linear | 30 | 512 | $30 \times 512 + 512 = 15,390$ |
| | Leaky ReLU | 512 | 512 | 0 |
| Fully Connected | Linear | 512 | 512 | $512 \times 512 + 512 = 262,656$ |
| | Leaky ReLU | 512 | 512 | 0 |
| Fully Connected | Linear | 512 | $5,000$ | $512 \times 5,000 + 5,000 = 2,560,512$ |

Supplementary Table 2: **Structure of scLAMBDA's decoder networks.**

| Layer | Detail | Input Size | Output Size | Number of Parameters |
|---|---|---|---|---|
| Fully Connected | Linear | 30×2 | 512 | $60 \times 512 + 512 = 30,780$ |
| | Leaky ReLU | 512 | 512 | 0 |
| Fully Connected | Linear | 512 | 512 | $512 \times 512 + 512 = 262,656$ |
| | Leaky ReLU | 512 | 512 | 0 |
| Fully Connected | Linear | 512 | 1 | $512 \times 1 + 1 = 513$ |

Supplementary Table 3: **Structure of scLAMBDA's mutual information estimator.**
